# Twinfilin1 controls lamellipodial protrusive activity and actin turnover during vertebrate gastrulation

**DOI:** 10.1101/2020.09.03.281659

**Authors:** Caitlin C. Devitt, Chanjae Lee, Rachael M. Cox, Ophelia Papoulas, José Alvarado, Edward M. Marcotte, John B. Wallingford

## Abstract

The dynamic control of the actin cytoskeleton is a key aspect of essentially all animal cell movements. Experiments in single migrating cells and *in vitro* systems have provided an exceptionally deep understanding of actin dynamics. However, we still know relatively little of how these systems are tuned in cell-type specific ways, for example in the context of collective cell movements that sculpt the early embryo. Here, we provide an analysis of the actin severing and depolymerization machinery during vertebrate gastrulation, with a focus on Twinfilin1. We confirm previous results on the role of Twf1 in lamellipodia and extend those findings by linking Twf1, actin turnover, and cell polarization required for convergent extension during vertebrate gastrulation.

## Introduction

Convergent extension (CE) is an evolutionarily conserved collective cell movement in which a group of cells interdigitates along a single axis, thus elongating the tissue in the perpendicular axis (Fig. 1A). CE drives the elongation of the body axis in essentially all animals and is critical for the shaping of diverse organs, including the kidney, heart, and cochlea, so defects in CE are directly implicated in human congenital anomalies, including neural tube closure defects and limb differences (Huebner and Wallingford, 2018; Shindo, 2018; Tada and Heisenberg, 2012; Wallingford et al., 2002).

**Figure 1.**
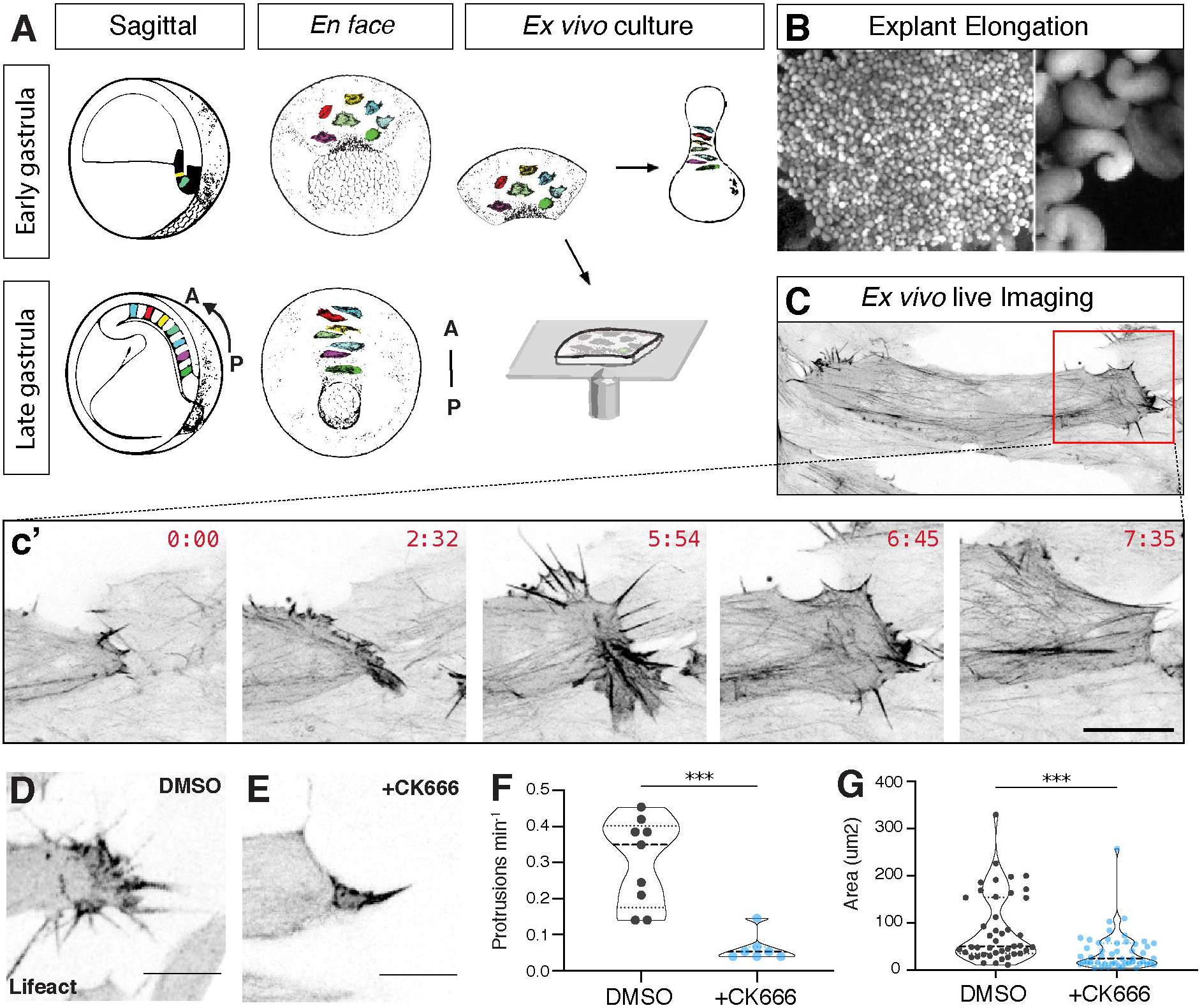
Lamellipodia in the *Xenopus* DMZ. **A)** Schematic showing CE cell movements and DMZ explant method. Cells are unpolarized at the start of gastrulation and over time take on a bipolar polarity and intercalate to form a longer, narrower array. **B)** Image showing elongated DMZ explants, close-up is shown at right. **C)** Still from time-lapse confocal microscopy showing a single round of lamellipodial extension and retraction from the mediolateral end of a single DMZ cell. **D)** DMSO does not affect DMZ lamellipodia. **E)** Typical effect of CK666 on DMZ lamellipodia. **F)** Quantification of effects of CK666 on protrusion frequency. DMSO vs CK666, p=0.0002**, n= 9 DMSO protrusions from 4 embryos, n=7 CK666 protrusions from 3 embryos. **G)** Quantification of effects of CK666 on protrusion size. DMSO vs CK666, p<0.0001***, n= 43 DMSO-treated protrusions from 4 embryos, n=49 CK666-treated protrusions from 3 embryos. Scale = 10um.

Recent work has demonstrated that CE in a wide array of cell types and organisms is driven by surprisingly similar cell biological mechanisms. From epithelial cells in mice and *Drosophila* to mesenchymal cells in amphibians, CE is achieved through the cooperative action of lamellipodia-based cell crawling and polarized junction contraction (reviewed in (Huebner and Wallingford, 2018; Shindo, 2018)). Despite its importance in development and disease, CE has received far less attention than other types of cell motility. For example, the integration of actin assembly and disassembly, actomyosin contraction, and cell adhesion have been exhaustively defined in single migrating cells (e.g. (Devreotes et al., 2017; Gardel et al., 2010; Lawson and Ridley, 2018)), while such systems remain only rudimentarily described in CE.

This gap in knowledge is significant, because while lamellipodia have been studied to immense depth in mammalian cultured cells, the extent to which those findings inform our understanding of CE remains undetermined. A very telling example relates to the classic work on Rho GTPases from Nobes and Hall. That central paper in cell biology linked Cdc42, Rac, and Rho to filipodia, lamellipodia, and formation of focal adhesions and stress fibers in migrating fiibroblasts (Nobes and Hall, 1995). However, though Rho and Rac are essential for CE in both *Xenopus* and mice (Habas et al., 2003; Ybot-Gonzalez et al., 2007), their functions do *not* appear to be well conserved during this collective cell movement. In the *Xenopus* DMZ during CE, Rac controls filipodia, while Rho and Rac cooperate to assembly lamellipodia, and Rho plays a distinct role in the control of cell shape and protrusion lifespan (Tahinci and Symes, 2003). Moreover, the roles for these GTPases in controlling lamellipodial activity differ even in sub-populations of mesodermal cells that engage in distinct types of collective cell movement in vertebrate embryos (Ren et al., 2006). Thus, further study of lamellipodia in the context of CE is warranted.

An especially intriguing question relates to actin assembly and turnover, two aspects of lamellipodial action that have been particularly well studied in migrating cells *in vitro* but have not been studied during CE. In single cells, actin filaments are polymerized at the leading edge by the interaction of membrane bound proteins, actin regulators, actin monomers, and existing actin filaments. As actin monomers are added to the distal end, existing filaments push on the plasma membrane, displacing the filament towards the cell-proximal portion on the lamellipodia in a process termed retrograde flow. At the rear of the lamellipodia, the lamella, focal adhesions attach actin fibers to the substrate or an extracellular matrix. Here, lamellipodial actin filaments are disassembled through filament severing and/or depolymerization (Carlier et al., 2015; Rottner and Schaks, 2019). The roles of actin regulatory proteins in migration of single cells have been documented extensively, including those of actin assembly proteins such as the Arp2/3 complex, and disassembly proteins such as Cofilin and Twinfilin (Kanellos and Frame, 2016; Poukkula et al., 2011; Swaney and Li, 2016). By contrast, we still know very little about how this machinery controls the dynamics of actin turnover or cell behavior during collective cell movement *in vivo*.

Interestingly, despite their centrality, mutation of core actin regulators in vertebrates frequently results in surprisingly specific developmental phenotypes. For example, Cofilin null mutants display defects in morphogenesis during neural tube closure in zebrafish and mice (Lin et al., 2010; Mahaffey et al., 2013), and the cofilin interactor cyclase-associated protein (Cap) is required for morphogenetic cell movements during gastrulation in zebrafish (Daggett et al., 2004; Daggett et al., 2007). Though the functionally related protein Twinfilin is required for a variety of actin-related cell behaviors in *Drosophila* (Wahlström et al., 2001; Wang et al., 2010), its role in the dynamic control of morphogenetic cell movement in developing vertebrates has not been defined.

Here, using the *Xenopus* dorsal gastrula mesoderm, we explored the localization dynamics of several actin-regulatory proteins, defined the *in vivo* interactome of Cofilin2, and explored the function of Twinfilin1 in convergent extension. These data provide an important *in vivo*, developmental complement to the extensive body of biochemical and cell biological work on Twinfilins and provide new insights into the role of core actin regulators in cell-type specific cell behaviors during vertebrate development.

## Results and Discussion

We chose to study lamellipodial protrusions in the dorsal gastrula mesoderm of amphibians (the so-called dorsal marginal zone; DMZ), an embryonic tissue that was among the first shown to undergo autonomous CE. In the 1940s, Holtfreter and Schectman independently showed that isolated explants of this tissue robustly elongate in culture (Holtfreter, 1944; Schectman, 1942), and pioneering work from Ray Keller and colleagues using *Xenopus* provided the first cell biological insights into this elongation process (Keller and Hardin, 1987; Shih and Keller, 1992a; Wilson and Keller, 1991). Since then, the tissue has served as a central paradigm for understanding the cellular and molecular basis of CE, including several molecular studies of lamellipodial protrusive activity (e.g. (Kim and Davidson, 2011; Pfister et al., 2016; Skoglund et al., 2008; Tahinci and Symes, 2003; Wallingford et al., 2000)).

These protrusions differ along the deep-superficial axis of the DMZ, and because our interest lay in comparing lamelliform protrusions in the *Xenopus* DMZ with the far more well-characterized lamellipodia in cultured cells, we performed a series of cell biological analyses focusing exclusively on the superficial lamelliform protrusions. We manually isolated DMZ explants and cultured them *ex vivo* for live imaging (Fig. 1A). We used a combination of confocal and TIRF imaging to monitor lamellipodial dynamics, protein localization, and actin dynamics.

In cultured cells, lamellipodia are formed by branched actin networks that are assembled by the Arp2/3 complex, so we first tested the role of this complex in the *Xenopus* DMZ. We found that treatment of DMZs with the Arp2/3 inhibitor Ck666 (Nolen et al., 2009) robustly suppressed the formation of lamelliform protrusions (Fig. 1D, E, F), and also dramatically altered the shape of those that did form. As a simple metric, we noted that lamellipodial area was significantly reduced after CK666 treatment (Fig. 1D, E, G).

We next quantified the localization patterns of several proteins (Fig. 2A, C, D) and compared these to the established patterns in lamellipodia of single cells in culture. We found that a PIP3 sensor (PH domain fused to GFP; (Tall et al., 2000)) was robustly enriched throughout the lamellipodia (Fig. 2C), similar to that described in cultured migrating cells. By contrast, the Myosin heavy chain Myh9 was very strongly enriched specifically in the proximal lamellipodium (Fig. 2D), again similar to that described for single migrating cells.

**Figure 2.**
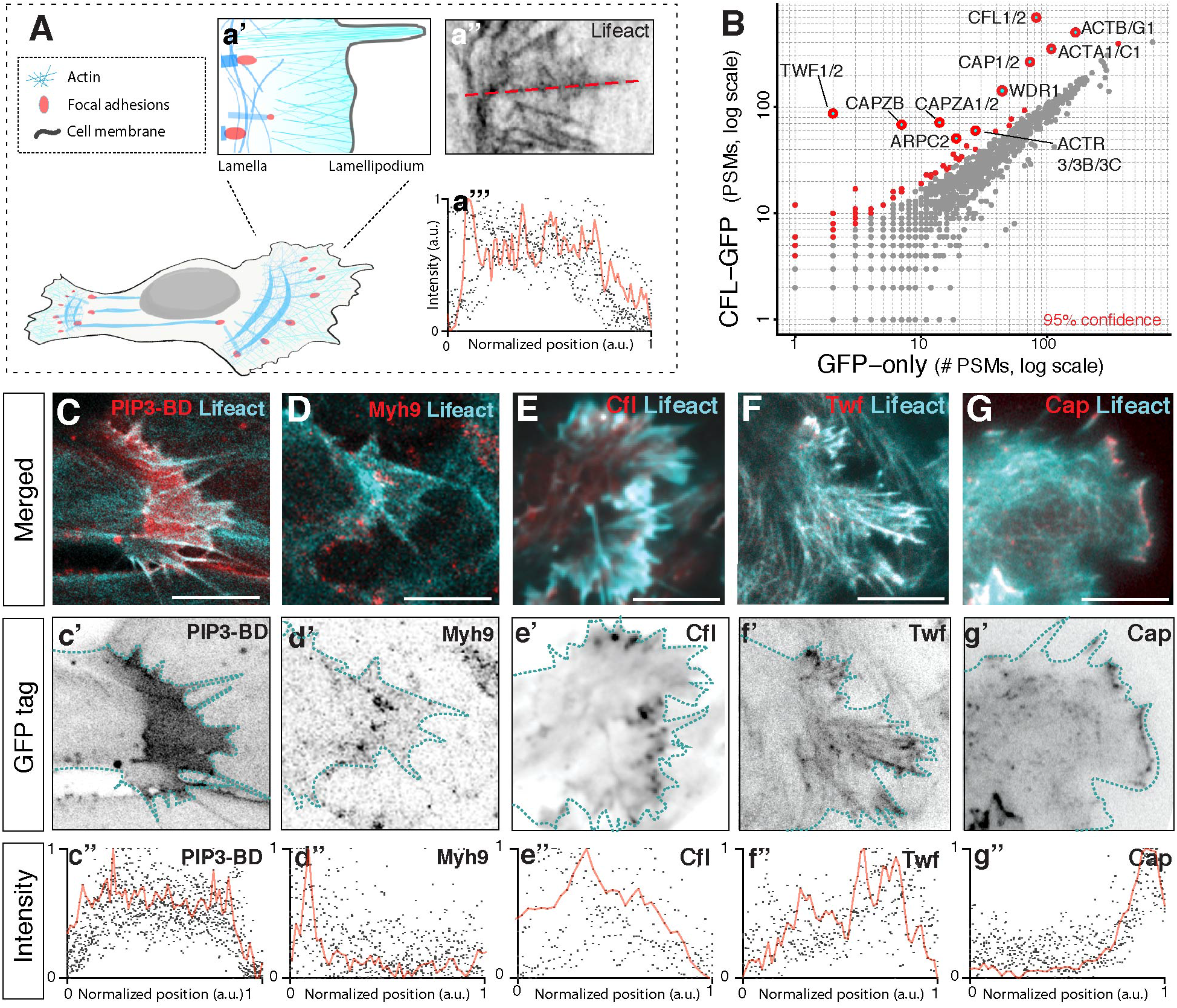
Localization of lamellipodial markers. **A)** Schematic showing the quantification scheme used here. **a”** Red dotted line indicates line-plot measurement taken along protrusions length. **a”’** Trace of actin intensity along protrusion length. Red line highlighting one example trace, black dots representative of several line plots. **B)** Interaction partners of Cfl2 were identified based on their enrichment in APMS of the Cfl2-GFP-tagged bait protein (vertical axes) relative to APMS of the untagged GFP controls (horizontal axes). Confidence values were calculated by one-sided Z-test (see Methods). A pseudocount of 1 PSM was added to each protein for visualization on a log-log plot. **C)** Pip3 localization and quantification; LifeAct in the alternate channel reports actin localization. **D)** Myh9 localization and quantification. **E)** Localization and quantification of Cfl2; LifeAct in the alternate channel reports actin localization. **F)** Localization and quantification of Twf1. **G)** Localization and quantification of Cap2. Scale= 10um

We next sought to explore the protein interaction landscape of actin regulation specifically in the *Xenopus* DMZ. We focused this experiment on Cofilin, which has been shown to be essential for CE in both mice and fish (Lin et al., 2010; Mahaffey et al., 2013). To identify interaction partners specifically in cells normally engaged in CE, we expressed GFP-tagged Cofilin2 by mRNA injection and manually dissected ~750 Keller explants and cultured them until NF stage 14 Fig. 1A, B). Protein was isolated from these tissue explants and affinity purification mass spectrometry (APMS) was performed using an anti-GFP antibody. To control for non-specific interactions, each experiment was accompanied by a parallel APMS experiment using un-fused GFP, the results of which were then used to subtract background by calculating a Z-test and fold change for differential enrichment for each protein (see Methods). Cofilin itself was by far the most strongly enriched protein recovered, providing a critical control for the specificity of this approach (Fig. 2B, Supp. Fig. 1; Supp. Table 1)

Our APMS data revealed that our APMS for Cofilin in the *Xenopus* DMZ was highly enriched for alpha-, beta-, and gamma-actin (Fig. 2B; Supp. Fig. 1; Supp Table 1), as expected and consistent with previous high-throughput protein interaction mapping in human cells (Rolland et al., 2014). Also enriched was the Cofilin regulator, Cyclase-Associated Protein (Cap)(Fig. 2B; Supp Table 1), consistent with data from cultured human cells and yeast (Moriyama and Yahara, 2002; Quintero-Monzon et al., 2009). We also identified significant, but less robust enrichment for subunits of the capping protein complex, as well as components of the Arp2/3 complex (Fig. 2B; Supp Table 1). Finally, we also observed highly significant enrichment for Twinfillin (Twf). Of course, APMS cannot distinguish between direct and indirect interactions, and it is very likely the case that the enrichment observed here APMS reflect association of all of these proteins with actin itself. Nonetheless, this experiment confirms in the vertebrate gastrula tight functional interactions that are known to link these proteins in yeast, in mammalian cultured cells, and *in vitro* (e.g. (Chaudhry et al., 2013; Goode et al., 1998; Hakala et al., 2019; Iwasa and Mullins, 2007; Johnston et al., 2015; Shekhar et al., 2019)).

We next used expression of GFP fusions to determine the localization of Cofilin and its two strongest interactors in our APMS data, Twf and Cap. We found that that Cofilin was broadly enriched in lamellipodia of mesoderm cells in the *Xenopus* DMZ, with a slight bias for the proximal region (Fig. 2E). Twf was also broadly enriched but with a slightly more distal bias (Fig. 2F). By contrast, Cap strongly was restricted specifically at the distal edge of the lamellipodium in these cells (Fig. 2G).

We then selected Twf for more in depth studies because while loss of either Cap or Cofilin in vertebrate are known to impact embryonic morphogenesis (Gurniak et al., 2005; Lin et al., 2010; Mahaffey et al., 2013) (Daggett et al., 2004; Daggett et al., 2007), the function of Twf in early vertebrate embryos has not been reported. We used morpholino-oligonucleotides to disrupt splicing of the Twf1 transcript, and knockdown of Twf1 elicited a robust defect in the elongation of the embryonic axis (Fig 3A; Supp. Fig. 2A-C). This defect was significantly rescued by re-introduction of Twf1 mRNA (Fig. 3A; Supp. Fig. 2C). As an additional control, we performed F0 knockout of Twf1 using CRISPR/Cas9, and this orthogonal loss of function approach elicited an identical phenotype (Fig. 3B; Supp. Fig. 2G). Together, these orthogonal loss-of-function approaches indicate that defective axis elongation is a specific result of Twf1 loss in *Xenopus*.

**Figure 3.**
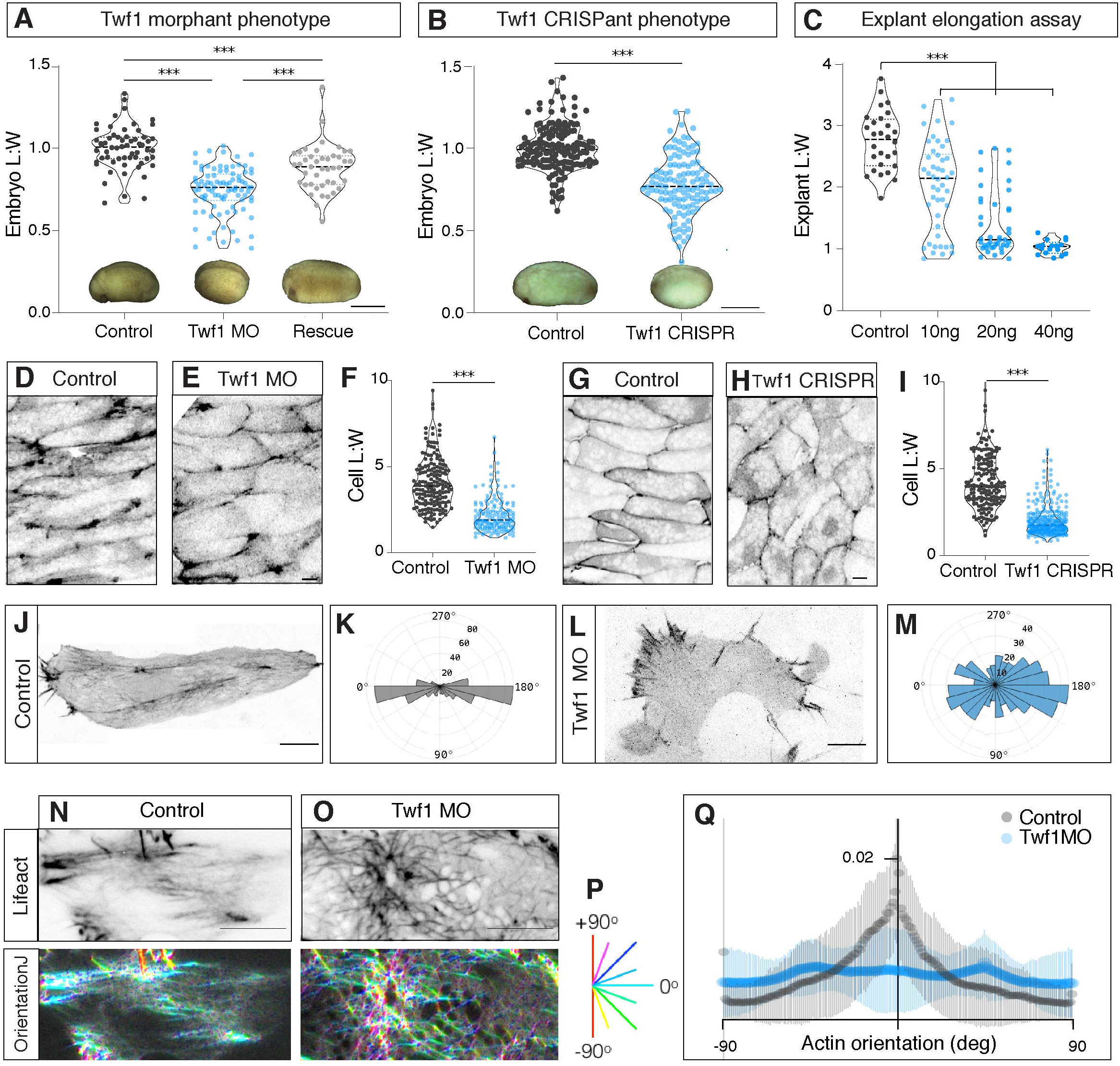
Twf1 is required for axis elongation and convergent extension. **A)** Quantification of axis elongation (as length/width ratio) for control, Twf1 morphants, and Twf1 mRNA rescued embryos (inset images show representative examples; see more embryos in Supp. Fig. 3C). Control vs Twf1 morphant, control vs rescue, Twf1 morphant vs rescue, each one-way ANOVA p<0.0001*** n= 60 control embryos, n=88 morphant embryos, n= 43 rescue embryos from 3 experiments. Scale = 1mm **B)** Quantification of axis elongation for control and Twf1 crispants, (inset images show representative examples). SgRNA only vs Twf1 Crispr, p<0.0001*** n= 155 control, n=119 Twf1 Crispr from 3 experiments. Scale = 1mm **C)** Quantification of explant elongation (as length/width ratio) for isolated DMZs from control and Twf1 morphant embryos. Control vs 10ng, 20ng, 40ng, p<0.0001*** in each one-way ANOVA, 20ng vs 40ns, n.s., (p=0.0966). n= 28 control, n=46 10ng, n= 41 20ng, n=17 40ng. **D, E)** Cell shapes for control and Twf1 morphant DMZs, as quantified in **F.** Control vs Twf1 morphant, p<0.0001*** n= 168 control cells from 6 embryos, n=156 Twf1 morphant cells from 6 embryos, 3 experiments. **G, H)** Cell shapes for control and Twf1 crispant DMZs, as quantified in **I**. Control vs Twf1 Crispr, p<0.0001*** n= 169 control cells from 11 embryos, n=294 Twf1 Crispr cells from 14 embryos, 3 experiments**. J)** Confocal image of mosaically labelled cell in control DMZ. **K)** Rose diagram showing the normal mediolateral polarization of lamellipodia in the control DMZ. **L)** Confocal image of mosaically labelled cell in Twf1 morphant DMZ. **M)** Rose diagram showing the disrupted mediolateral polarization of lamellipodia in the morphant. **K,M)** Control vs Twf1 morphant, k.s. tests p<0.0001*** n= 119 control protrusions from 5 embryos, n=194 Twf1 morphant protrusions from 9 embryos, 3 experiments. **N, O)** TIRF images of actin cables in control and Twf1 morphant DMZ cells, in lower panels, color reflects orientation of cables as indicated by the legend in **P. Q)** Quantification of actin cable organization in control and Twf1 morphants. DMSO vs CK666, p<0.0001*** n= 6 control embryos, n=11 Twf1 morphant embryos. Scale= 10um, unless otherwise noted.

Curiously, the robust elongation defect in embryos lacking Twf1 was not associated with the severe dorsal flexion commonly observed following disruption of convergent extension by manipulations of PCP signaling (e.g. (Wallingford and Harland, 2001)). Nonetheless, Twf1 KD also elicits a dose-dependent suppression of the elongation of isolated DMZ explants (Fig. 3C; Supp. Fig. 2E, F), which is known to be driven by convergent extension in a tissue-autonomous manner (Keller et al., 1992). Moreover, disruption of Twf1 either by MO or by CRISPR resulted in a significant reduction of cellular length-to-width ratios in the DMZ (Fig. 4D-I), a phenotype that is commonly associated with defective CE (e.g. (Tahinci and Symes, 2003; Wallingford et al., 2000)). Together these data demonstrate that Twf1 is required for normal convergent extension in *Xenopus*.

**Figure 4.**
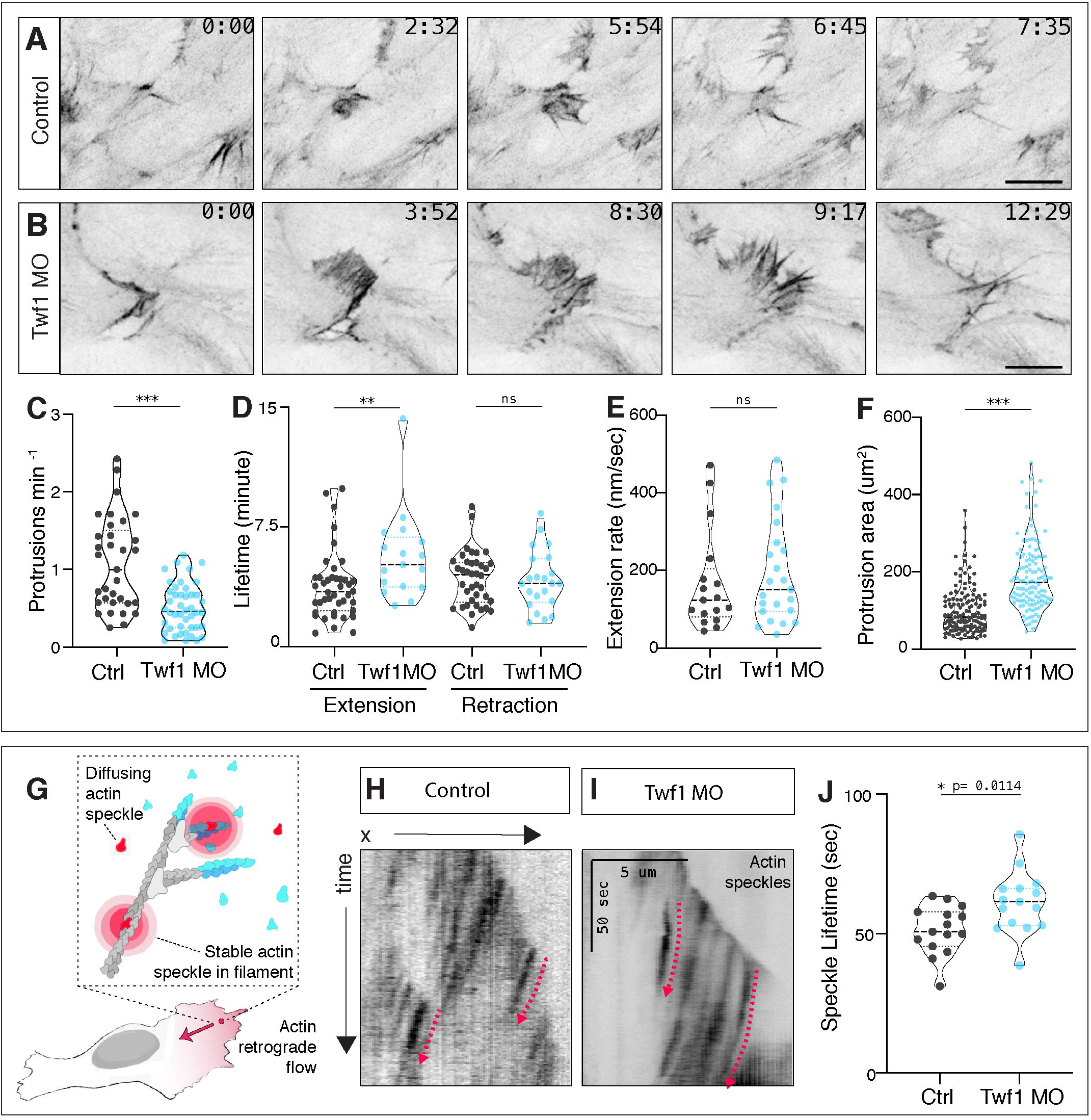
Twf1 controls lamellipodial dynamics in the *Xenopus* DMZ. **A, B)** Confocal images of single cells in a LifeAct-GFP expressing control and Twf1 morphant DMZs. Scale= 10um. **C)** Quantification of lamellipodial protrusion frequency for control and Twf1 morphants. Control vs Twf1 morphant, p<0.0001*** n= 33 control protrusions from 2 embryos DMSO, n= 44 Twf1 morphant protrusions from 4 embryos. **D)** Quantification of lamellipodial extension and retraction lifetimes. Control vs Twf1 morphant, extension lifetime p=0.0096**, Control vs Twf1 morphant retraction lifetime, n.s. n= 42 control protrusions from 4 embryos, n= 22 Twf1 morphant protrusions from 6 embryos. **E)** Quantification of lamellipodial extension rate. Control vs Twf1 morphant, n.s., p=0.5294. n= 17 control protrusions from 4 embryos, n=23 Twf1 morphant protrusions from 5 embryos. **D)** Quantification of lamellipodial area for control and Twf1 morphants. Control vs Twf1 morphant, p<0.0001***. n= 144 control protrusions from 3 embryos, n=127 Twf1 morphant protrusions from 6 embryos. **G)** Schematic of fluorescent speckle imaging. **H, I)** TIRF images showing fluorescent actin speckles in lamellipodia in control and Twf1 morphant *Xenopus* DMZs. Scale (x, distance)= 5um, scale (y, time)= 50 seconds. **G)** Quantification of actin speckle lifetimes. Control vs Twf1 morphant, p=0.0114*. n= 15 control protrusions from 3 embryos, n=15 Twf1 morphant protrusions from 6 embryos.

To understand the cell biological basis for this defect, we asked how Twf1 loss impacted the robust mediolateral polarization of actin and lamellipodia in DMZ cells during convergent extension. While lamellipodia were tightly confined to mediolateral cell faces in control DMZ explants, as expected (Shih and Keller, 1992a), this mediolaterally polarity was lost when Twf1 was disrupted (Fig. 3J-M). This loss of the mediolateral polarity of lamellipodia is commonly associated with defective CE, for example by disruption of PCP or Rho signaling (Tahinci and Symes, 2003; Wallingford et al., 2000).

The function of lamellipodia in driving cell movement remains poorly understood in CE, as compared to the context of individual migrating cells in culture. For example, cells during CE are bipolar and frequently form lamellipodia on both ends, preventing the concept of a “leading edge” to be used in this context (Shih and Keller, 1992a). Moreover, lamellipodia are thought to collaborate with junction contractions to cooperatively drive the movement of cell bodies during CE (Huebner and Wallingford, 2018). Finally, a specialized cytoplasmic “node-and cable” system has been described in *Xenopus* DMZ cells (Kim and Davidson, 2011; Skoglund et al., 2008). How this system relates to the cytoskeleton in the cell body of single migrating cells (i.e. stress fibers, etc.) remains unknown, and the dynamics of the DMZ node and cable system may integrate lamellipodial activity and junction contraction (Shindo et al., 2019). We therefore used TIRF imaging to ask how Twf1 loss impacts the node and cable system in the DMZ.

In normal cells, these actin cables are strongly aligned in the mediolateral axis of the cell (Kim and Davidson, 2011; Skoglund et al., 2008). Using the Orientation J plugin in Fiji (Püspöki et al., 2016), we observed a similar polarity of actin cables, as expected (Fig. 3N, P, Q). By contrast, this polarization was significantly disrupted by Twf1 loss, with cables displaying a near random orientation (Fig. 3O, P, Q). Interestingly, we did not observe Twf1 localization to the actin cables themselves (not shown), consistent with the absence of Twf1 at mammalian stress fibers (Vartiainen et al., 2000). These results suggest that the defects in actin cable polarity are likely secondary to the defects in lamellipodial position or dynamics.

To better understand how Twf1 function in lamellipodia may impact cell polarization during CE, we next sought to quantify the effects of Twf1 loss on the dynamic behavior of lamellipodia. This is an important un-answered question, actually, because while extensively studied in biochemical terms, we still know little about the role of Twinfilins in living cells. For example, loss of Twf1 in *Drosophila* S2 cells in culture results in enlarged lamellipodia with altered actin dynamics, and similar results were recently reported in B16-F1 mouse melanoma cells (Hakala et al., 2019; Iwasa and Mullins, 2007). There are as of yet no reports of the impact of Twf1 loss on lamellipodial dynamics in an intact tissue *in vivo*.

We therefore used targeted microinjection to generate mosaic embryos in which cells labelled with LifeAct-GFP were surrounded by unlabeled cells, allowing unambiguous quantification of lamellipodial activity in the DMZ. Consistent with the reports from cell culture, Twf1 loss in the *Xenopus* DMZ did not impact the overt morphology of lamellipodia (Fig. 4A, B), but did severely disrupt the *dynamics* of lamellipodial assembly: Twf1 KD cells initiated lamellipodial extension less frequently (Fig. 4C). Moreover, though extension rates were not altered, lamellipodia spent relatively more time elongating compared to controls (Fig. 4D, E). Lamellipodia retraction times were not different between control and Twf1 KD (Fig. 4E), and accordingly, the mean size of lamellipodia in these cells was significantly larger (Fig. 4F).

Finally, we sought to link the extensive previous biochemical characterization of Twf to the *in vivo* functions we describe here. In assays using single actin filaments, the mechanisms by which Twf and its associated factors tune severing and depolymerization have been extensively characterized (e.g. (Hakala et al., 2019; Hilton et al., 2018; Johnston et al., 2015)). However, the subcellular concentration of these proteins is unknown *in vivo*, and the rates of actin turnover vary considerably between *in vitro* studies and those in living cells (Miyoshi and Watanabe, 2013). To quantify actin turnover *in vivo* specifically during *Xenopus* CE and to assess the impact of Twf1 loss, we developed methods to perform actin fluorescent speckle microscopy on DMZ explants (Fig 4G). We injected fluorescently labeled actin protein into 4-cell stage *Xenopus* embryos, and imaged DMZ explants using TIRF microscopy. We titrated the dose of injected actin until we achieved appropriate distribution of speckles, which in our hands was 0.1ng injected per blastomere.

We found that actin speckle lifetimes were significantly increased by Twf1 disruption in *Xenopus* gastrula mesoderm (Fig 4 H, I, J), consistent with actin speckle data from *Drosophila* S2 cells (Iwasa and Mullins, 2007). Our result is also consistent with recent FRAP data on actin turnover in mouse melanoma cells (Hakala et al., 2019). Together with those previous reports, our findings argue that the cellular functions of Twf1 in both lamellipodial dynamics and actin turnover are evolutionarily conserved in *Drosophila* (Iwasa and Mullins, 2007), mice (Hakala et al., 2019), and *Xenopus*. This result is of interest because while Twf1 is among the very few actin-regulatory proteins that have been conserved in evolution from protozoa to vertebrates (De Melo et al., 2008), yeast and mouse Twinfilins differ substantially in their activities when assayed biochemically (Hilton et al., 2018). Further *in vivo* characterization of Twf1, as well as related factors such as Cap and Cofilin therefore should be illuminating.

## Conclusions

Here we show that many elements of lamellipodia-related protein localization and proteinprotein interactions defined in single cells in culture are conserved in cells of the *Xenopus* DMZ during CE. More importantly, we provide here the first dynamic analysis of Twinfilin function on cell behaviors in a vertebrate *in vivo*, showing that loss of Twf1 results in defects in the dynamics of lamellipodia shape and lifetime, as well as in defects in actin turnover. While on the one hand, these results confirm *in vivo* results obtained previously in cell culture, on the other it is important because both the upstream regulation and the downstream function of lamellipodia during CE are highly specialized. For example, we find that DMZ cells lacking Twf1 function fail to elongate mediolaterally (Fig. 3), a phenotype also observed with disruption of either PCP or Rho signaling, but not Rac signaling (Tahinci and Symes, 2003; Wallingford et al., 2000), potentially suggesting links between the former proteins and Twf1. Moreover, how force for cell body displacement is achieved during CE remains a contentious issue (Huebner and Wallingford, 2018), so our finding that specific disruption of lamellipodial dynamics following Twf1 disruption is associated with failure to polarize cytoplasmic actin cables in the DMZ is also significant. Asking how Twf1 loss impacts junction contraction in these cells will be an interesting next step. Ultimately, these data advance our goal of an integrated view of the mechanisms linking ubiquitous and evolutionarily conserved machinery of actin regulation to the highly specialized behaviors of diverse cell types in vertebrate animals.

## Acknowledgements

We thank Shashank Shekhar for discussions and critical comments on the manuscript and Anna Battenhouse for advice on proteomic analyses. This work was supported by the NICHD (R01HD099191) and the NIGMS (R01GM104853). E.M.M. also acknowledges support from the NIH (R35GM122480) and Welch Foundation (F-1515).

## Experimental Procedures

Please see the Supplemental Experimental Procedures. All animal experiments were approved by the IACUC of the University of Texas at Austin, protocol no. AUP-2012-00156.

## Author Contributions

C.D. and J.W. conceived the project. C.D., C.L., J.W. and E.M. designed experiments. C.D. performed all imaging and analysis. C.L. and O.P. performed the APMS experiments, and R.C. and E.M. analyzed APMS data. J.W. and C.D. wrote the paper with input and editing from all authors.

## Supplemental Experimental Procedures

### *Xenopus* embryo manipulations

*Xenopus laevis* females were super-ovulated by injection of hCG (human chorionic gonadotropin). The following day, eggs were squeezed from the females. *In vitro* fertilization was performed by homogenizing a small part of a harvested testis in 1X Marc’s Modified Ringer’s (MMR) and mixing with collected eggs. Embryos were dejellied in 3% cysteine (pH 7.9) at the two-cell stage. Fertilized embryos were rinsed and reared in 1/3X Marc’s Modified Ringer’s (MMR) solution. For microinjections, embryos were placed in a 2% ficoll in 1/3X MMR solution and injected using forceps and an Oxford universal micromanipulator. Morpholino oligonucleotide or CRISPR/Cas9 was injected into two dorsal blastomeres to target the dorsal marginal zone (DMZ). Embryos were microinjected with mRNA at the 4-cell stage for uniform labeling and at the 32-to 64-cell stage for mosaic labeling. Embryos were staged according to Nieuwkoop and Faber (Nieuwkoop and Faber, 1994).

### Plasmids, mRNA, protein, and MOs for microinjections

*Xenopus* gene sequences were obtained from Xenbase (www.xenbase.org) and open reading frames (ORF) of genes were amplified from the *Xenopus* cDNA library by polymerase chain reaction (PCR), and then are inserted into a pCS10R vector fused with C-terminal GFP. The following constructs were cloned into pCS vector: Twf1-GFP, Cap1-GFP, Cfl2-GFP. These constructs were linearized and the capped mRNAs were synthesized using mMESSAGE mMACHINE SP6 transcription kit (ThermoFisher Scientific). Concentration for GFP localization was titrated to lowest concentration where we could still detect GFP signal, but no cellular phenotype was observed. The amount of injected mRNAs per blastomere are as follows: Lifeact-GFP or –RFP [50-100pg], PIP3-PBD [25pg] (Tall et al.), Myl9-GFP [20pg] (Shindo and Wallingford, 2014), Twf1-GFP [25pg for imaging, 1ng for rescue], Cap1-GFP[25pg], and Cfl2 [25pg].

Twf1 morpholino was designed to target exon-intron splicing junction (https://www.gene-tools.com/). The MO sequences and the working concentrations include: MO: 5’-TGAGTCAAAACACTTACATGGGAGT-3’, 10 ng per injection unless otherwise noted.

CRISPR sgRNAs were designed to target exon 2 using the online design tool CHOPCHOP (Labun et al., 2019). Synthetic sgRNA was ordered from Synthego (https://www.synthego.com/products/crispr-kits/synthetic-sgrna) and combined with Cas9 protein (PNABIO, Cat no CP01) prior to injection at 4-cell stage. 2 individual targets were tested in isolation (250pg/10nl injection) and pooled (125pg each), each giving the same embryonic phenotype. Pooled sgRNAs were used for all experiments. The sgRNA sequences and working concentrations include: Twf1 Target 1: ACCAAAUUCCUUCUUCACAGUGG (125pg/10nl injection), Twf1 Target 2: AAAAUGCGCAAGGCUUCGAGUGG (125pg/10nl injection), Cas9 protein (1ng/10nl injection).

For fluorescent actin speckle microscopy, labeled actin monomers (Actin568, ThermoFisher, Catalog number A12374) were reconstituted in a 2mg/ml stock solution. A 0.01 ug/ul working solution was prepared for each 10nl injection, and 0.1ng was injected into dorsal blastomeres of 4-cell stage embryos. This concentration was determined empirically by titrating the amount injected until no cell phenotype was observed and labeled actin monomers were sparse enough to detect individual “speckles” (Watanabe and Mitchison, 2002).

### RT-PCR

To verify the efficiency of Twf1 MOs, MOs were injected into all cells at the 4-cell stage and total RNA was isolated using the TRIZOL reagent (Invitrogen) at stage 14. cDNA was synthesized using M-MLV Reverse Transcriptase (Invitrogen) and random hexamers. Twf1 cDNAs were amplified by Taq polymerase (NEB) with these primers: Twf1F 5’-ACA CCA GAC ACT TCA AGG AAT AG-3’, Twf1R 5’-GGT TCT TGC AGC TGG AGT ATA A-3’, ODC 426F 5’-GGC AAG GAA TCA CCC GAA TG-3’ and ODC 843R 5’-GGC AAC ATA GTA TCT CCC AGG CTC-3’.

### Embryo phenotyping and Keller explant elongation assay

To determine the morpholino oligonucleotide concentration to use, we used a dose curve and chose the lowest concentration of morpholino while maintaining consistent phenotype, deciphered based on embryo and explant phenotypes (Fig 3 supp). Embryo and Keller explant length and width measurements were taken in Fiji, as previously described (Shih and Keller, 1992b; Wallingford et al., 2000).

### Live imaging of Keller explants

mRNAs were injected at the dorsal side of 4 cell stage embryos, and the dorsal marginal zone (DMZ) tissues were dissected out at stage 10.5 using forceps, hairloops, and eyebrow hair knives. Each explant was mounted to a fibronectin-coated dish in Steinberg’s solution (Shindo and Wallingford, 2014), and cultured at 16°C for half a day before imaging with a Zeiss LSM700 confocal microscope or Nikon N-STORM combined TIRF/STORM microscope.

### Image analysis

Images were processed with the Fiji distribution of ImageJ and Photoshop (Adobe) software suites, and figures were assembled in Illustrator (Adobe). Cell length and width measurements and protrusion angles were manually taken in Fiji along the long axis and through the centroid of the cell. To define the area of individual protrusions, custom scripts (available upon request) were written in Matlab to create a binary mask with uniform threshold, define individual protrusions, and count the number of pixels representing the area of individual protrusions. Actin cable orientations were taken from TIRF images using the OrientationJ plugin in Fiji (Püspöki et al., 2016) (http://bigwww.epfl.ch/demo/orientation/). Speckle tracking was performed using MosaicSuite (http://mosaic.mpi-cbg.de PMID: 16043363) object tracking plugin in Fiji. Custom scripts were written in Matlab to count the lifetime of each speckle, and then converted to msec based on imaging time interval. Statistical analyses were carried out using Prism (Graphpad) software with student t tests or one-way ANOVAs for significance.

### Immunoprecipitation from *Xenopus* DMZs for mass spectrometry

mRNA encoding GFP-only or Cfl2-GFP was injected into 2 dorsal blastomeres of the 4-cell stage *Xenopus* embryo. Approximately 1,000 DMZs per sample were isolated from stage 10.5 embryos using forceps and microblades and were cultured in 1X Steinberg’s solution (0.58 mM NaCl, 0.64 mM KCl, 0.33 mM Ca(NO2)2, 0.8 mM MgSO4, 5 mM Tris, 50 μg/ml gentamicin, pH 7.4-7.6) until stage-matched embryos reached stage 14. The cultured explants were collected and immunoprecipitation (IP) was performed using GFP-Trap Agarose Kit (ChromoTek, cat# gtak-20). Immunoprecipitated proteins were eluted in 2X sample buffer.

### Affinity purification-mass spectrometry

Immunoprecipitated proteins were resuspended in SDS-PAGE sample buffer and heated 5 minutes at 95°C before loading onto a 7.5% acrylamide mini-Protean TGX gel (BioRad). After 7 minutes of electrophoresis at 100 V, the gel was stained with Imperial Protein stain (Thermo) according to manufacturer’s instructions. The protein band was excised, diced to 1 mm cubes and processed by standard trypsin in-gel digest methods for mass spectrometry. Digested peptides were desalted with Hypersep Spin Tip C-18 columns (Thermo Scientific), dried, and resuspended in 60 μl of 5% acetonitrile, 0.1% acetic acid for mass spectrometry.

Mass spectrometry was performed on a Thermo Orbitrap Fusion Lumos Tribrid mass spectrometer, using a data-dependent acquisition strategy and four technical replicates per sample. Peptides were separated using a 60-minute gradient reverse phase chromatography method on a Dionex Ultimate 3000 RSLCnano UHPLC system (Thermo Scientific) with a C18 trap to Acclaim C18 PepMap RSLC column (Dionex; Thermo Scientific). Scans were acquired in rapid scan mode using a top-speed method with dynamic exclusion after 1 time for 20 seconds and stepped HCD collision energies of 27, 30 and 33%.

Raw MS/MS spectra were processed using Proteome Discoverer (v2.3). We used the Percolator node in Proteome Discoverer to assign unique peptide spectral matches (PSMs) at FDR < 5% to the composite form of the *X. laevis* reference proteome described in (Drew et al., 2020). In order to identify proteins statistically significantly associated with Cfl2, we calculated both a log2 fold-change and a Z-score for each protein based on the observed PSMs in the Cfl2-GFP experimental pulldown relative to the GFP-only control pulldown. We calculated significance for protein enrichment in the experiment relative to control using a one-sided Z-test as in (Lu et al., 2007) with a 95% confidence threshold (z ≥ 1.645). We observed high concordance between technical replicates (Supp. Fig 1).

### Data deposition

Proteomics data have been deposited into the MassIVE repository (accession MSV000086057) and ProteomeXchange (accession PXD021258).

---

**Supplemental Figure 1.**
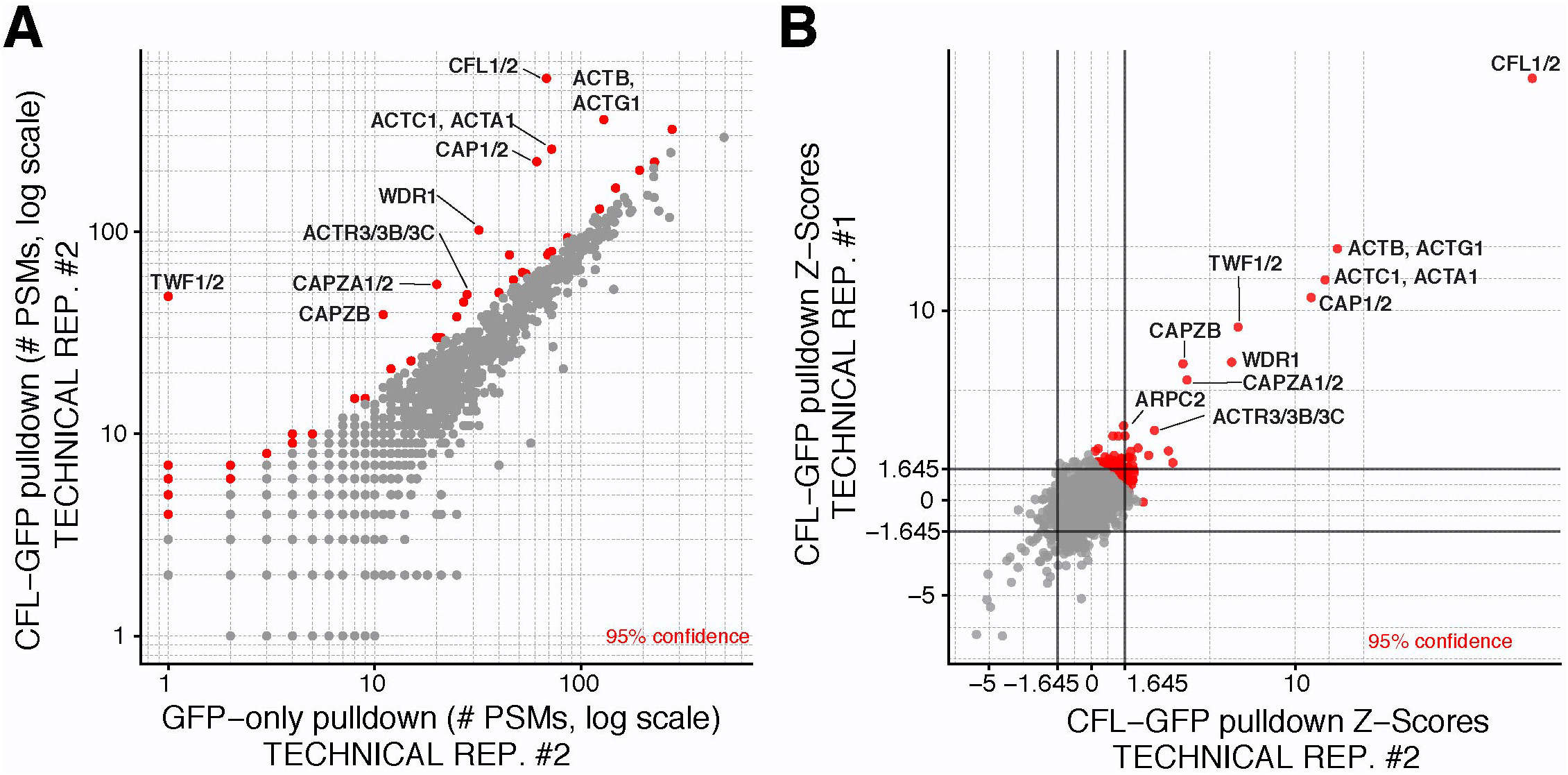
**A)** Enriched prey proteins after Cfl2-GFP pulldown were identified based on their enrichment in APMS (vertical axes) relative to APMS of the untagged GFP controls (horizontal axes). Confidence values were calculated by one-sided Z-test (see Methods). A pseudocount of 1 PSM was added to each protein for visualization on a log-log plot. **B)** Comparison of technical replicates (Panel A and Fig 2B). Confidence values are based on a joint Z-score, calculated by summing the Z-score for each protein in each technical replicate and dividing by the square root of *N* replicates (*N* = 2).

**Supplemental Figure 2.**
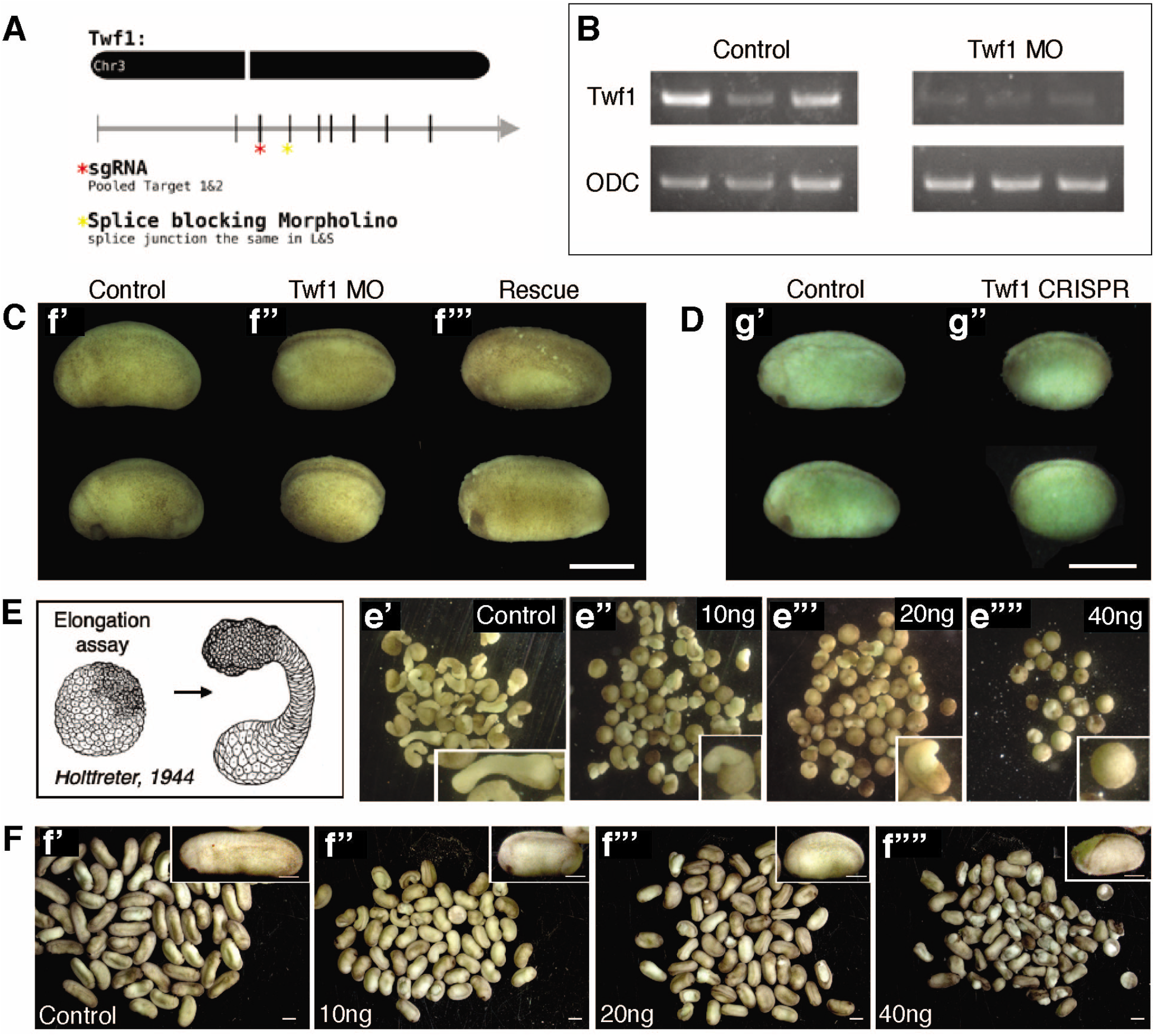
**A)** Schematic of Twf1 gene and our targeting reagents. **B)** RT-PCR showing disruption of Twf1 transcript splicing by the Twf1 MO. **C)** Representative embryos for Twf1 MO-mediated knockdown. **D)** Representative embryos for Twf1 CRISP-mediated knockdown. **E)** Schematic of DMZ explant elongation assay. **e’)** Twf1 morphant DMZs fail to elongate. **F’)** Twf1 morphant embryos fail to elongate.

**Supplemental Table 1:** This Excel spreadsheet contains all APMS data, ranked by Z-score confidence for 99th, 95th, and 90^th^ percentiles (see Methods). Note that Cofilin itself is by far the highest scoring protein, providing a key positive control.

